# Innovative Carnitine-Fed Rats Model Reveals Resveratrol Butyrate Ester’s Multimechanistic Role in Reducing TMAO and Cardiovascular Risk

**DOI:** 10.1101/2024.12.16.628813

**Authors:** Chih-Yao Hou, Cai-Sian Liu, Ming-Kuei Shih, Asif Ali Bhat, You-Lin Tain, Chang-Wei Hsieh, Yu-Wei Chen, Shin-Yu Chen

**Affiliations:** Department of Seafood Science, College of Hydrosphere, National Kaohsiung University of Science and Technology, Kaohsiung 81157, Taiwan; Graduate Institute of Food Culture and Innovation, National Kaohsiung University of Hospitality and Tourism, Kaohsiung 81271, Taiwan; Department of Pediatrics, Kaohsiung Chang Gung Memorial Hospital, Kaohsiung 83301, Taiwan; Institute for Translational Research in Biomedicine, Kaohsiung Chang Gung Memorial Hospital, Kaohsiung 83301, Taiwan; College of Medicine, Chang Gung University, Taoyuan 33305, Taiwan; Department of Medical Research, China Medical University Hospital, Taichung City 40402, Taiwan; Department of Food Science and Biotechnology, National Chung Hsing University, Taichung 40227, Taiwan; Department of Food Science, National Pingtung University of Science and Technology, Pingtung 91201, Taiwan

**Author notes:** These authors contributed equally to this work (Chih-Yao Hou, Cai-Sian Liu). **Correspondence:** Yu-Wei Chen, Department of Food Science and Biotechnology, National Chung Hsing University, 145 Xingda Rd., South Dist., Taichung City 402, Taiwan. Shin-Yu Chen, Department of Food Science, National Pingtung University of Science and Technology, 1 Shuefu Road, Neipu, Pingtung 91201, Taiwan.

**Keywords:** resveratrol butyrate ester, TMAO, carnitine, OCTN2, microbiota, metabolomics, cardiovascular protection, innovative animal model

## Abstract

**BACKGROUND:** Trimethylamine-N-oxide (TMAO), a metabolite produced from dietary carnitine through gut microbiota, is a recognized risk factor for cardiovascular disease (CVD). High-fat diets and carnitine supplementation exacerbate TMAO levels and related risks, making them valuable in developing experimental models for studying CVD. Resveratrol butyrate ester (RBE) and its monomer ED4 have shown potential in reducing TMAO levels and improving cardiovascular outcomes through metabolic and microbial modulation, but their precise mechanisms remain unclear.

**METHODS AND RESULTS:** A novel animal model was established using 36 male Sprague-Dawley rats fed a high-fat diet supplemented with carnitine to elevate serum TMAO levels, simulating dietary-induced CVD risks. Rats were divided into six groups: control diet (CN), high-fat diet (HFD), high-fat diet with carnitine (HFDC), HFDC with dimethylbutanol (HFDCB), CN with ED4 (CNM), and HFDC with ED4 (HFDCM). Physiological parameters, serum lipid profiles, SCFA levels, microbiota composition, and gene expression (OCTN2 and FMO3) were analyzed. ED4 supplementation reduced serum TMAO levels by upregulating OCTN2 expression, promoting urinary TMAO excretion, and restoring SCFA levels. ED4 also modulated gut microbiota, reducing TMA-producing bacteria (e.g., *Bacteroides*), and improved cardiovascular markers, including reduced blood lipid levels and fat accumulation. While dimethylbutanol inhibited FMO3 expression to reduce TMAO, ED4 acted primarily through carnitine utilization and microbiota modulation. Both treatments enhanced urinary TMAO excretion and altered gut microbiota composition.

**CONCLUSIONS:** This study introduced an innovative animal model combining high-fat and carnitine-enriched diets to study TMAO-related cardiovascular risks. ED4 demonstrated multimechanistic effects in reducing TMAO levels and CVD risk factors by modulating gut microbiota, restoring SCFA levels, and enhancing carnitine metabolism. These findings highlight ED4’s therapeutic potential in cardiovascular protection and metabolic regulation. However, further research is needed to elucidate the molecular pathways underlying ED4’s effects on TMAO metabolism and its broader translational applications.

## Introduction

Cardiovascular disease (CVD) has become a major global health issue due to its high mortality and morbidity rates.^1,2^ A key factor in CVD is atherosclerosis, a chronic inflammatory condition affecting the inner lining of arteries.^3^ Atherosclerosis typically arises from dyslipidemia or angiogenesis disorders,^4^ making CVD often characterized by dyslipidemia and elevated cholesterol levels.^5^ Furthermore, CVD is closely linked to dietary habits.^6^ In recent years, an increasing number of individuals have adopted high-calorie, high-fat diets,^7^ which are typically rich in both fat and carnitine. While carnitine is promoted as a dietary supplement to enhance athletic performance,^8^ its role in increasing the risk of CVD has also been highlighted.

Dietary carnitine is metabolized by the gut microbiota into trimethylamine (TMA),^9^ which is subsequently oxidized in the liver by flavin-containing monooxygenases (FMOs) to form trimethylamine N-oxide (TMAO).^10^ TMAO plays a critical role in the development of CVD, particularly in promoting atherosclerosis.^11^ As a result, dietary carnitine consumption leads to increased TMAO levels, thereby accelerating the progression of atherosclerosis.^12^ Furthermore, diets rich in fat and carnitine not only significantly increase TMAO production but also disrupt gut microbial homeostasis.^13,14^

Recent metagenomic sequencing studies have established links between the gut microbiota and numerous diseases, including CVD.^15^ Studies have shown that TMAO’s impact on gut flora serves as a significant marker for CVD in humans,^14,16^ highlighting the complex interplay between dietary patterns, gut microbiota, and cardiovascular health.^17^ Given the strong association between dietary habits and CVD,^6^ modifying the gut microbiota has emerged as one of the most effective strategies to lower TMAO levels in the bloodstream and reduce cardiovascular risk.^18^

Prebiotics have been shown to promote the growth of beneficial gut bacteria while reducing bacteria responsible for TMA production. For example, studies have demonstrated that resveratrol (RSV) can modify gut microbiota and decrease TMA production in mice.^15^ RSV can alter gut microbial composition, reverses dysbiosis induced by high-fat diets, and mitigates cardiovascular risk factors. Therefore, RSV has also been highlighted for its potential in inhibiting TMAO levels and exerting anti-CVD effects, which are attributed to its rapid metabolism and ability to modulate gut microbiota.^19,20^ Additionally, RSV’s modulation of gut microbiota influences energy metabolism and satiety hormones, contributing to its anti-obesity effects.^21^

However, RSV’s limited bioavailability presents a challenge.^22^ To address this, resveratrol butyrate ester (RBE) has been synthesized, offering improved water solubility, superior absorption, an extended half-life, and enhanced cardiovascular benefits compared to RSV.^23,24^ Previous co-culture studies have shown that both RSV and RBE promote the growth of *Bifidobacterium longum* while suppressing *Clostridium asparagiforme*, thereby reducing TMA production. Particularly, RBE has proven effective in reducing TMA levels.^25^ Furthermore, the monoester derivative 3-O-butanoylresveratrol (ED4) exhibits better anti-obesity effects compared to the diester form (ED2) in high-fat diet-induced obesity models.^26^ These findings underscore the promising therapeutic potential of RSV derivatives in cardiovascular and metabolic health.

Building on our previous findings that resveratrol butyrate ester (RBE) and its purified monomer exhibit significant potential in modulating gut microbiota, reducing TMA production, and preventing TMAO-induced cardiovascular disease (CVD), this study takes a comprehensive approach to evaluate the therapeutic efficacy of ED4. Given the urgent need for innovative experimental models to address the growing burden of CVD, we successfully established a novel animal model using a high-fat, carnitine-supplemented diet to induce elevated serum TMAO levels. This model serves as a robust platform for investigating the biological activities of ED4, including its capacity to mitigate obesity, regulate genes involved in TMA and TMAO metabolism, and modulate gut microbiota composition.

Specifically, this research aims to assess ED4’s effects on physiological parameters, liver and intestinal tissue expressions of FMO3 and OCTN2, and biochemical and microbiological profiles in serum and urine. By leveraging this innovative model, we seek to elucidate ED4’s mechanisms of action and its therapeutic potential in reducing CVD risk. This study provides valuable insights into ED4’s ability to modulate metabolic and microbial pathways and contributes to the development of novel strategies for combating TMAO-associated CVD.

## Methods and Materials

### Animals and Experimental Design

In this study, 36 male Sprague-Dawley (SD) rats (average body weight, 95 ± 4 g, age: 4 weeks) were obtained from BioLASCO Taiwan Co., Ltd. (Taipei, Taiwan). The experiments were approved by the Committee for Laboratory Animals of the National Kaohsiung University of Science and Technology (approval No.: 0109-AAAP-012).

The total experimental period was 8 weeks. After a 1-week housing period with regular diet (AIN-93G, Taipei, Taiwan), the rats were randomly assigned into different groups with experimental diets from week 1-8, including the control diet (CN), high fat diet (HFD), HFD with 0.5% carnitine diet (HFDC), HFDC with 1% DMB in drinking water (HFDCB), CN with ED4 (CNM), and HFDC with ED4 (HFDCM) groups (n=6). DMB, a structural analogue of choline, functions as a non-lethal inhibitor of TMA production by intestinal microbes in nutrient-rich environments. It holds promise as a therapeutic agent for CVD by specifically targeting microbial TMA production, thereby reducing circulating TMAO levels. The efficacy of DMB in suppressing TMA production has been demonstrated in mice fed a high-choline diet.^27,28^ In this study, DMB is utilized for cross-comparison with RBE to explore the relationship between carnitine metabolism and gut microbiota.

The experiment design and feed composition are shown in supplementary Fig S.1 and Table S.1. The animals in the HFDCB was treated with 1% DMB in the drinking water from week 3-8, while the CNM and HFDCM groups were treated by daily oral gavage of 20 mg/kg BW of ED4 from weeks 3-8. The provided dose of ED4 was based on a previous study, administered at 20 mg/kg BW/day.^29^ Daily food intake and consumption were recorded throughout the study. Prior to sacrifice, blood pressure was measured, and fecal and urine samples were collected for further analysis. Blood pressure data was obtained using the indirect tail-cuff technique (BP-2010 Series).

In week 8, following a 10-hour fasting period, rats were administered an intramuscular anesthetic cocktail consisting of a 1:1 combination of Zoletil (25 mg/kg) and Rompun (23.32 mg xylazine hydrochloride/kg). Blood samples were then collected via cardiac puncture. Organs were harvested and weighed, with the liver, retroperitoneal fat, and aortic arch tissues preserved in 10% formalin for staining. The remaining tissues were quickly frozen in liquid nitrogen and stored at -80°C for subsequent analyses.

### Serum Biochemical Indicator and Tissue Histopathology

The blood samples were centrifuged at 3000×g for 10 min at 4°C, and the supernatant was collected to obtain the serum samples stored at -80°C until analyses. In this study, the triglycerides (TG), total cholesterol (TC), and high-density lipoprotein cholesterol (HDL-C) contents were analyzed by an automated clinical chemistry analyzer (NX-500i, Fuji, Masaaki Takei, Japan). The atherosclerosis index (AI) was calculated using the formula (TC – HDL-C) / HDL-C.

After being fixed in 10% formalin, the liver and retroperitoneal fat were embedded in paraffin, baked at 65°C for 60 minutes, sectioned, and then stained using the H&E technique. The stained sections were imaged using the MoticEasyScan system and analyzed with the Motic DS Assistant (4K). The aortic arch was cryo-embedded in OCT gel and then sectioned. After staining with Oil Red O, the slides were mounted using a water-based gel and analyzed with ImageJ 1.50i (National Institutes of Health, USA). The coronary artery area to be analyzed was circled, and the color analysis method was applied to calculate the positive staining area, which was then expressed as a percentage of the total coronary artery area.

### Short-Chain Fatty Acid

The pretreatment of serum and fecal samples and the GC/MS analysis conditions were performed following the method described by Zhang et al.^30^

### Microbiota Analysis

The extraction and treatment of fecal samples, as well as the next-generation sequencing and bioinformatic analysis of the gut microbiota, were carried out following the method described by Chen et al.^26^

### Serum DMA, TMA and TMAO Analysis

Serum samples were spiked with the internal standard diethylamine and separated using HPLC (Agilent 1200, California, USA). The concentrations of dimethylamine (DMA), TMA, and TMAO in the serum were then analyzed using LC-MS/MS (Agilent 6410, California, USA) equipped with an electrospray ionization source. Separation was performed on a SeQuant ZIC-HILIC column (150 × 2.1 mm, 5 µm; Merck KGaA, Darmstadt, Germany). The mobile phase consisted of 20% methanol with 15 mmol/L ammonium formate and 80% acetonitrile, at a flow rate of 0.3 to 1 mL/min.

### Gene Protein Analysis

OCTN2 protein was extracted from rat jejunum using the Rat Solute Carrier Family 22 Member 5 (SLC22A5) ELISA Kit. The extraction and operational procedures followed the kit instructions, and absorbance at 450 nm was measured using an ELISA reader.

FMO3 protein expression in rat liver was analyzed by first treating the liver tissue with 250 µL of RIPA lysis buffer and zirconium beads. Homogenization was performed on ice, and the supernatant was collected after centrifugation. The protein concentration was adjusted to 5 µg/µL, with an equal amount of sample buffer added, followed by heating in a dry bath at 95°C for 10 minutes. The samples were then separated by SDS-PAGE and transferred onto a PVDF membrane. The membrane was blocked in 5% skim milk at room temperature, shaking at 50 rpm for 1 hour. Following blocking, the membrane was incubated with the primary antibody for FMO3 at 4°C with shaking for 12 hours. After removing the primary antibody, the membrane was washed three times with buffer, then incubated with the secondary antibody at room temperature for 10 minutes with shaking. After removing the secondary antibody, the membrane was washed three more times with buffer, and protein bands were visualized using chemiluminescent detection under a luminescence imaging system.

### Urine Metabolomics

Urine samples were filtered using Amicon 3,000 molecular weight (MW) cutoff filters. The filtrate was collected, and 65 µL of the internal standard, 4.67 mM DSS-D6 (Chenomx Inc., Edmonton, Alberta, Canada), was added. PBS was subsequently added to bring the total volume to 650 µL, with the exact volume recorded. The pH of each sample was adjusted to 6.8 by adding small amounts of 1 N HCl or NaOH. Proton NMR spectra were acquired using a 600 MHz NMR spectrometer (JEOL ECZ600R, Tokyo, Japan). Metabolite identification and quantification were carried out using Chenomx NMRSuite 9.02 (Chenomx Inc., Edmonton, Canada).^31^ Metabolite concentrations in the samples were normalized based on 24-hour urine collection records.

### Statistics

Results are expressed as means ± standard errors (S.E.). One-way analysis of variance was applied as appropriate. Duncan’s multiple range test was performed as a post hoc multiple comparison test. The significance differences between examined groups were set at *P < 0.05*. All statistical analyses were performed using the Statistical Analysis System software package (SAS Institute, Cary, NC, USA).

## Results

In this study, DMB is utilized for cross-comparison with RBE to explore the relationship between carnitine metabolism and gut microbiota.

### Physiological Characteristics

This study aimed to investigate the association between TMAO precursors (such as carnitine), CVD-related parameters, therapeutic compounds (ED4), and gut microbiota. A high-fat, carnitine-enriched diet was used to induce TMAO production in rats, a recognized risk factor for CVD. The effects of ED4 supplementation were compared with those of DMB treatment and evaluated based on growth characteristics, as presented in Table 1. In the CNM group, which received ED4 supplementation, the visceral fat percentage (2.17%) was significantly lower than that of the CN group (2.74%), with no other significant differences observed. These findings suggest that ED4 supplementation did not produce adverse effects in the rats.

**Table 1.** Effect of ED4 on physiological characteristics of rats with cardiovascular disease induced by high-fat complex carnitine diet.

|  | CN | HFD | HFDC | HFDCB | CNM | HFDCM |
| --- | --- | --- | --- | --- | --- | --- |
| Food intake (g) | 33.17±11.42 | 32.21±12.36 | 32.03±8.89 | 26.93±11.52 | 29.78±15.05 | 29.03±9.49 |
| Body weight (g) | 376.67±8.40 | 517.13±23.73* | 478.73±17.86* | 412.97±20.09# | 338.63±9.49<sup>#</sup> | 414.23±13.43 |
| Weight gain (%) | 58.09±0.77 | 75.21±1.36* | 70.72±0.36* | 66.39±1.06* | 55.01±1.58<sup>#</sup> | 60.58±1.37<sup>#</sup> |
| Liver weight (%) | 3.35±0.17 | 2.67±0.16 | 2.88±0.16* | 3.07±0.11# | 3.14±0.20<sup>#</sup> | 3.27±0.14<sup>#</sup> |
| Kidney weight (%) | 0.82±0.02 | 0.69±0.02 | 0.71±0.04 | 0.93±0.00*<sup>#</sup> | 0.78±0.04 | 0.77±0.03 |
| Visceral fat (%) | 2.74±0.22 | 4.23±0.18* | 3.96±0.23* | 2.65±0.24# | 2.17±0.07<sup>*#</sup> | 4.37±0.15 |
| Retroperitoneal fat weight (%) | 1.57±0.12 | 2.33±0.15 | 1.97±0.28 | 1.20±0.37 | 1.30±0.13 | 2.33±0.17 |
| Epididymal fat weight (%) | 1.15±0.09 | 2.04±0.04* | 1.79±0.14* | 1.19±0.07# | 1.02±0.10<sup>#</sup> | 1.99±0.08 |
| SBP (mmHg)<sup>a</sup> | 124.73±2.15 | 139.15±4.45 | 136.45±4.87 | 119.95±3.63# | 112.57 ±5.87<sup>#</sup> | 114.67±8.01 |
| DBP (mmHg)<sup>b</sup> | 90.65±1.22 | 106.18±4.89 | 101.63±3.85 | 92.30±6.35 | 78.77±3.41 | 78.03±9.22 |
| TG (mg/dL)<sup>c</sup> | 63.67±2.85 | 76.33±5.7 | 86.33±7.86 | 37.67±0.33<sup>#</sup> | 43.33±10.67<sup>#</sup> | 49.67±1.45<sup>#</sup> |
| TC (mg/dL)<sup>d</sup> | 41.00±3.51 | 56.00±2.65* | 60.67±5.21* | 47.00±2.08 | 45.33±1.86<sup>#</sup> | 49.67±2.03 |
| HDL (mg/dL)<sup>e</sup> | 29.00±3.46 | 30.67±2.33 | 33.33±2.19 | 46.00±5.51 | 34.00±4.16 | 46.33±2.73* |
| AI<sup>f</sup> | 0.43±0.13 | 0.39±0.05 | 0.63±0.06 | 0.29±0.04<sup>#</sup> | 0.30±0.06 | 0.23±0.11 |

The data are presented as mean ± standard error of each group. \* $p < 0.05$ vs. CN; # $p < 0.05$ vs. HFDC.
CN: Normal diet, normal water; HFD: High-fat diet, normal water; HFDC: High-fat diet with 0.5% carnitine diet, normal water; HFDCB: High-fat diet with 0.5% carnitine diet, 1% DMB in drinking water; CNM: Normal diet, normal water, tube feeding 20 mg/Kg/Day ED4; HFDCM: High-fat diet with 0.5% carnitine diet, normal water, tube feeding 20 mg/Kg/Day ED4.
<sup>a</sup>Systolic blood pressure.
<sup>b</sup>Diastolic blood pressure.
<sup>c</sup>Triglycerides.
<sup>d</sup>Total cholesterol.
<sup>e</sup>High-density lipoprotein cholesterol.
<sup>f</sup>Atherosclerosis index.

The HFD and HFDC groups showed significantly higher weight gain, visceral fat, and epididymal fat compared to the CN group, despite no significant differences in dietary intake. However, the HFDC group exhibited slightly lower weight gain and fat mass than the HFD group, indicating that while both a high-fat diet and a high-fat-carnitine diet lead to increased body weight and fat content in rats, the addition of carnitine may slightly mitigate these effects.

For blood pressure measurements, both systolic (SBP) and diastolic blood pressure (DBP) were slightly elevated in the HFD and HFDC groups compared to the CN group, indicating a trend toward increased blood pressure with high-fat or high-fat-carnitine diets. However, no significant differences were noted between the HFD and HFDC groups.

In terms of blood biochemical markers, TG and TC levels were elevated in both the HFD and HFDC groups compared to the CN group. Specifically, TG and TC levels in the HFD group were 1.2 and 1.37 times higher than in the CN group, while in the HFDC group, TG and TC were 1.36 and 1.48 times higher. These findings suggest that the high-fat-carnitine diet led to a greater increase in TG and TC levels than the high-fat diet alone. Additionally, the AI in the HFDC group was 0.63, which was higher than the CN group’s 0.43 and the HFD group’s 0.39. This suggests that the addition of carnitine to the diet may further contribute to an increase in the atherosclerosis index (AI).

The HFDCB group experienced a weight gain of 66.39%, slightly lower than the 70.72% observed in the HFDC group. The percentages of visceral fat and epididymal fat were 2.65% and 1.19%, respectively, significantly lower than the 3.96% and 1.79% in the HFDC group. Although the retroperitoneal fat percentage was also reduced, this difference was not statistically significant. Blood pressure and lipid analysis showed that both SBP and DBP were lower in the HFDCB group compared to the HFDC group. Specifically, SBP was 119.95 mmHg in the HFDCB group, significantly lower than the 136.45 mmHg recorded in the HFDC group. TG levels in the HFDCB group were 37.25 mg/dL, which is 0.43 times that of the HFDC group, indicating a significant difference. Additionally, lower levels of TC and a reduced AI were observed. These results suggest that adding DMB to the drinking water effectively reduces fat weight, blood pressure, and blood lipid levels in rats fed a high-fat, carnitine-enriched diet, though its effect on overall body weight was relatively modest.

In the HFDCM group, body weight increased by 60.58%, significantly lower than the 70.72% observed in the HFDC group. However, the percentages of visceral fat and epididymal fat were not significantly different from those in the HFDC group, suggesting that administration of the purified ED4 monomer effectively reduced body weight gain associated with the high-fat, carnitine-enriched diet, but did not significantly affect body fat percentage. Additionally, blood pressure measurements showed that the SBP and DBP in the HFDCM group were 114.67 and 78.03 mmHg, respectively, which were lower than the 136.45 and 101.63 mmHg observed in the HFDC group. Although these reductions were not statistically significant, they indicate a potential role of ED4 in regulating blood pressure. Blood lipid analysis revealed that TG levels in the HFDCM group were 49.67 mg/dL, 0.57 times lower than those in the HFDC group. Furthermore, ED4 supplementation showed a tendency to reduce TC and AI, indicating its potential to lower blood lipid levels.

### Histological Analysis

The results of the histological analysis are presented in Fig. 1. The HFD and HFDC groups demonstrated significant liver vacuolization compared to the CN group (Fig. 1a). The fat droplet areas in both the HFD and HFDC groups were significantly larger than those in the CN group, with the HFDC group showing slightly larger fat droplets than the HFD group. The mesh structure area of retroperitoneal fat in the HFD group was significantly greater than in the CN group, while the HFDC group had a significantly smaller area than both the CN and HFD groups. As shown in Table 1, HFD treatment effectively induced visceral and subcutaneous fat accumulation in the rats. In contrast, the HFDC group did not have a significant effect on visceral fat accumulation but significantly inhibited subcutaneous fat accumulation.

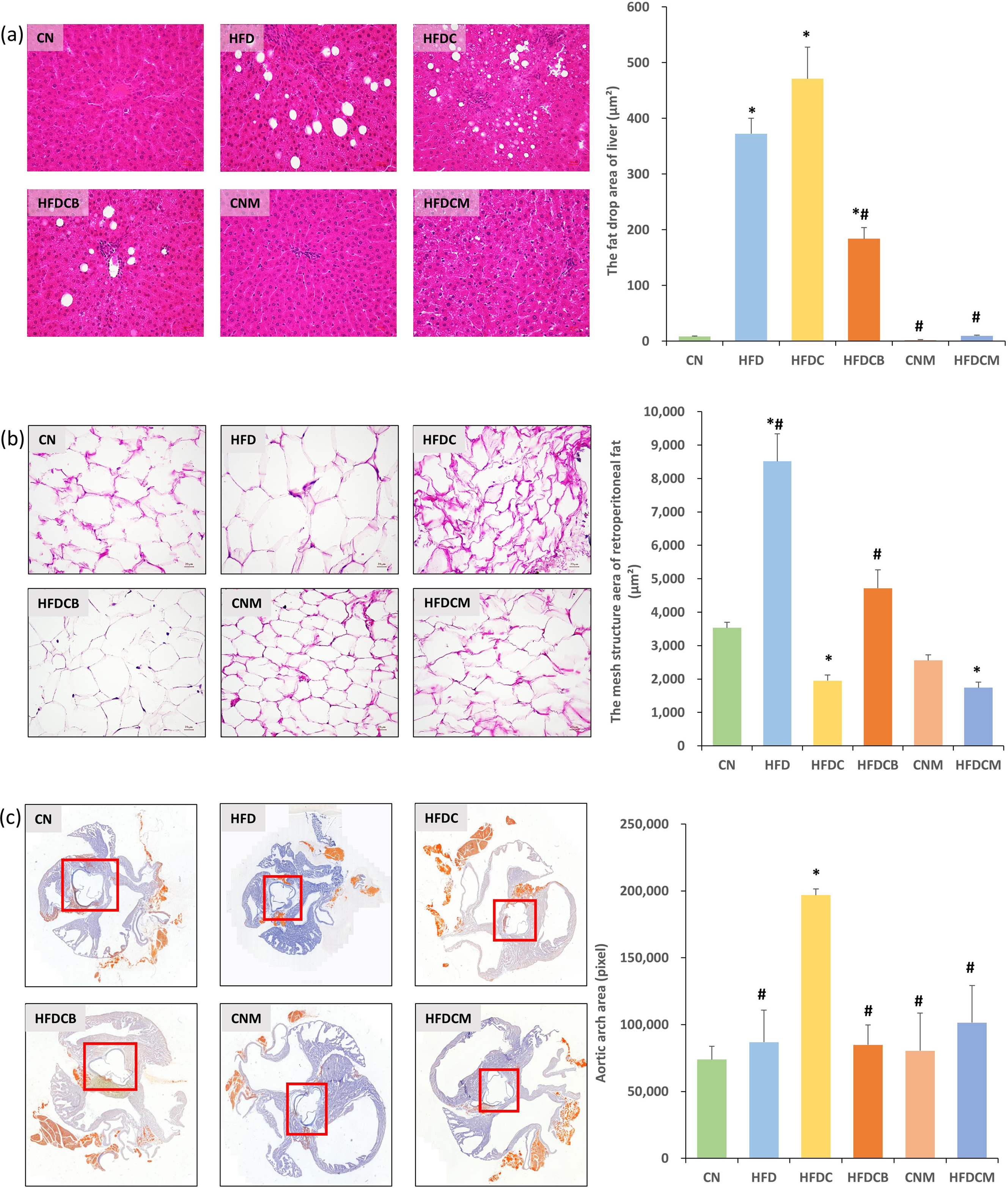

The fat droplet areas in the livers of the HFDCB and HFDCM groups were significantly lower than those in the HFDC group (Fig. 1a), suggesting that supplementation with DMB and ED4 effectively reduced liver fat accumulation. However, the mesh structure areas of retroperitoneal fat in the HFDCB and HFDCM groups were not significantly different from those in the HFDC group (Fig. 1b). Interestingly, the fat vacuole area in the HFDCB group was actually higher than that observed in the HFDC group.

The AI values in the HFDCB and HFDCM groups were both lower than those in the HFDC group (Fig. 1c). These findings indicate that while carnitine increases the atherosclerosis risk index, both DMB and ED4 supplementation reduce this index, highlighting their potential to lower the risk of atherosclerosis.

### SCFA Analysis

This study evaluates SCFA content in the body by measuring acetic acid, propionic acid, and butyric acid levels in rat serum and feces (Table 2). The results show that serum SCFA levels in the HFD and HFDC groups were lower than those in the CN group. However, in the HFDCB and HFDCM groups, serum SCFA levels were higher than in the HFDC group. Notably, the concentrations of acetic acid and propionic acid in these groups returned to levels comparable to the CN group after treatment with DMB and ED4. In terms of fecal SCFA levels, the concentrations of acetic acid and propionic acid were lower in the HFD and HFDC groups compared to the CN group. In contrast, the HFDCB and HFDCM groups showed significantly higher levels of fecal acetic acid and propionic acid than the HFDC group, with both differences reaching statistical significance. However, these concentrations remained not significantly different from those in the CN group. These findings suggest that a high-fat diet decreases SCFA content in rats, while treatment with DMB and ED4 leads to an upward trend in SCFA levels.

**Table 2.**
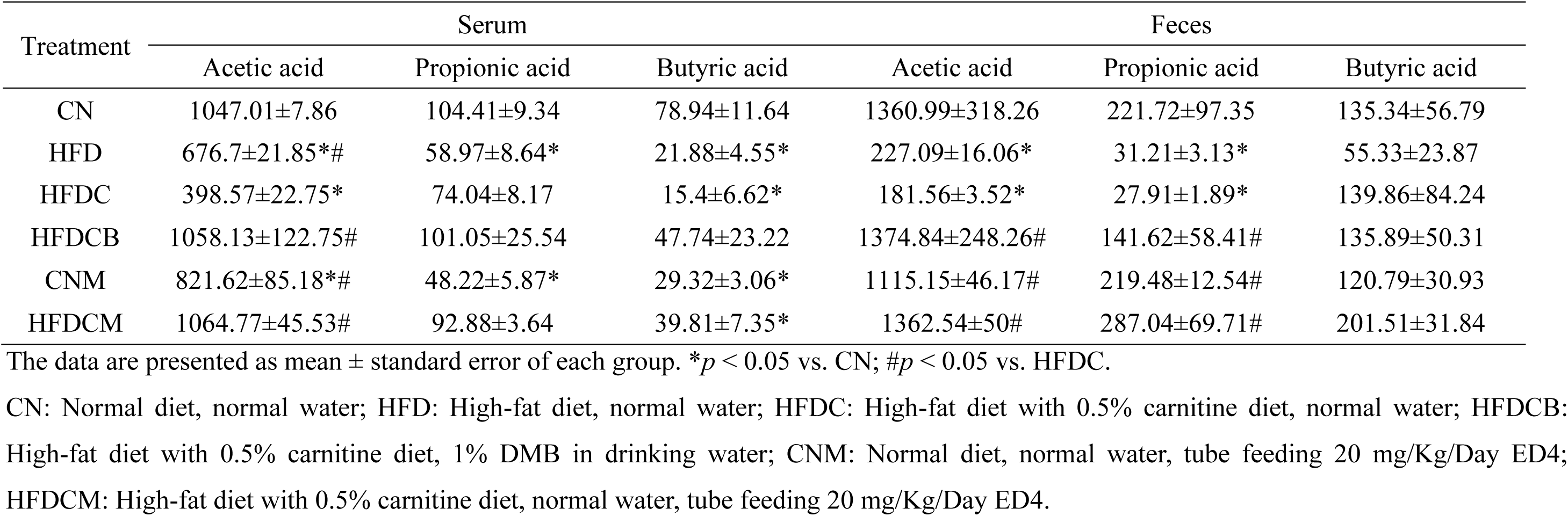
Effect of ED4 on serum and fecal short-chain fatty acid levels in rats with cardiovascular disease induced by high-fat complex rnitine diet.

| Treatment | Serum |  |  | Feces |  |  |
| --- | --- | --- | --- | --- | --- | --- |
|  | Acetic acid | Propionic acid | Butyric acid | Acetic acid | Propionic acid | Butyric acid |
| CN | 1047.01±7.86 | 104.41±9.34 | 78.94±11.64 | 1360.99±318.26 | 221.72±97.35 | 135.34±56.79 |
| HFD | 676.7±21.85*# | 58.97±8.64* | 21.88±4.55* | 227.09±16.06* | 31.21±3.13* | 55.33±23.87 |
| HFDC | 398.57±22.75* | 74.04±8.17 | 15.4±6.62* | 181.56±3.52* | 27.91±1.89* | 139.86±84.24 |
| HFDCB | 1058.13±122.75# | 101.05±25.54 | 47.74±23.22 | 1374.84±248.26# | 141.62±58.41# | 135.89±50.31 |
| CNM | 821.62±85.18*# | 48.22±5.87* | 29.32±3.06* | 1115.15±46.17# | 219.48±12.54# | 120.79±30.93 |
| HFDCM | 1064.77±45.53# | 92.88±3.64 | 39.81±7.35* | 1362.54±50# | 287.04±69.71# | 201.51±31.84 |

The data are presented as mean ± standard error of each group. \* $p < 0.05$ vs. CN; # $p < 0.05$ vs. HFDC.
CN: Normal diet, normal water; HFD: High-fat diet, normal water; HFDC: High-fat diet with 0.5% carnitine diet, normal water; HFDCB: High-fat diet with 0.5% carnitine diet, 1% DMB in drinking water; CNM: Normal diet, normal water, tube feeding 20 mg/Kg/Day ED4; HFDCM: High-fat diet with 0.5% carnitine diet, normal water, tube feeding 20 mg/Kg/Day ED4.

### Serum DMA, TMA, and TMAO

The analysis of DMA, TMA, and TMAO levels in rat serum showed no significant differences between the HFD and CN groups (Fig. 2), indicating that seven weeks of high-fat diet treatment does not significantly alter serum DMA or TMA levels in rats, although a slight increase in TMAO was observed. In contrast, the HFDC group, which received additional dietary carnitine, had serum concentrations of DMA, TMA, and TMAO at 124.00, 118.00, and 4031.33 ng/mL, which were significantly higher than the 85.67, 3.67, and 70.00 ng/mL observed in the HFD group. This suggests that carnitine supplementation increases DMA, TMA, and TMAO levels by approximately 1.4, 32, and 58 times, respectively, compared to the high-fat diet alone, demonstrating that adding carnitine to a high-fat diet significantly elevates serum concentrations of TMA-related metabolites.

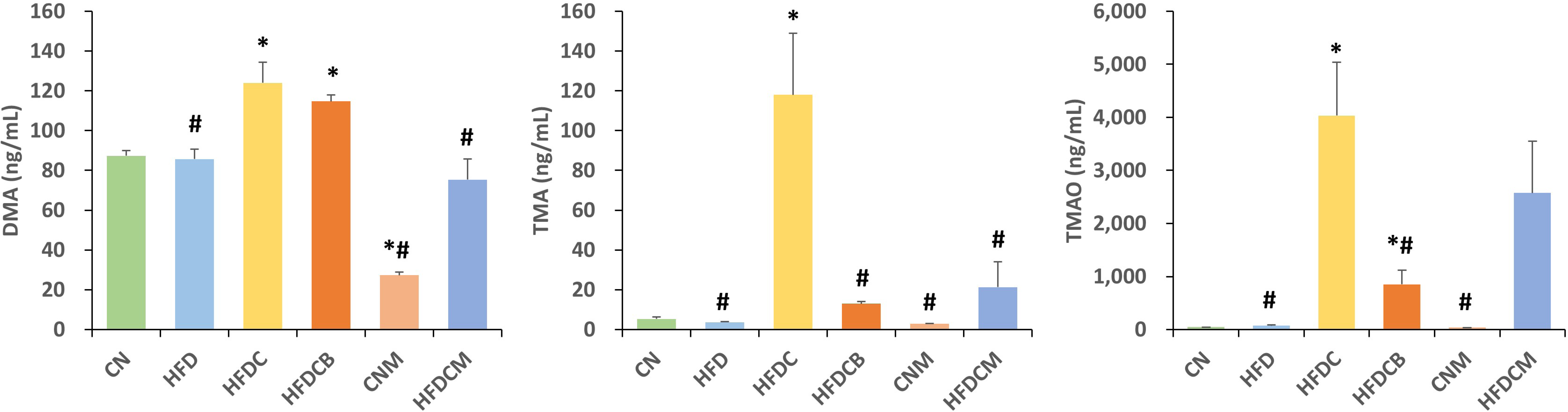

In the HFDCB group, serum TMA and TMAO levels were 13.00 and 849.67 ng/mL, representing reductions of 9.1 and 4.7 times compared to the HFDC group. Similarly, in the ED4-treated HFDCM group, TMA and TMAO levels were 21.33 and 2575.33 ng/mL, showing reductions of 5.5 and 1.6 times, respectively, compared to the HFDC group. These results demonstrate that both DMB and ED4 treatments effectively lower serum TMA and TMAO levels.

### Urine Metabolites

Urine metabolomics analysis using quantitative NMR revealed seventeen detected metabolites (Table 3). In the PCA plot (Fig. 3a), the groups fed an HFD mixed with carnitine were separated from the others along PC2 (10.3%). The treatment groups, HFDCB and HFDCM, formed a distinct cluster and were separated from the HFDC group along PC1 (80.2%), suggesting that the urine metabolite profiles of HFDCB and HFDCM are highly similar. Furthermore, HFDCB and HFDCM exhibited higher metabolite levels than the HFDC group (Table 3).

**Table 3.**
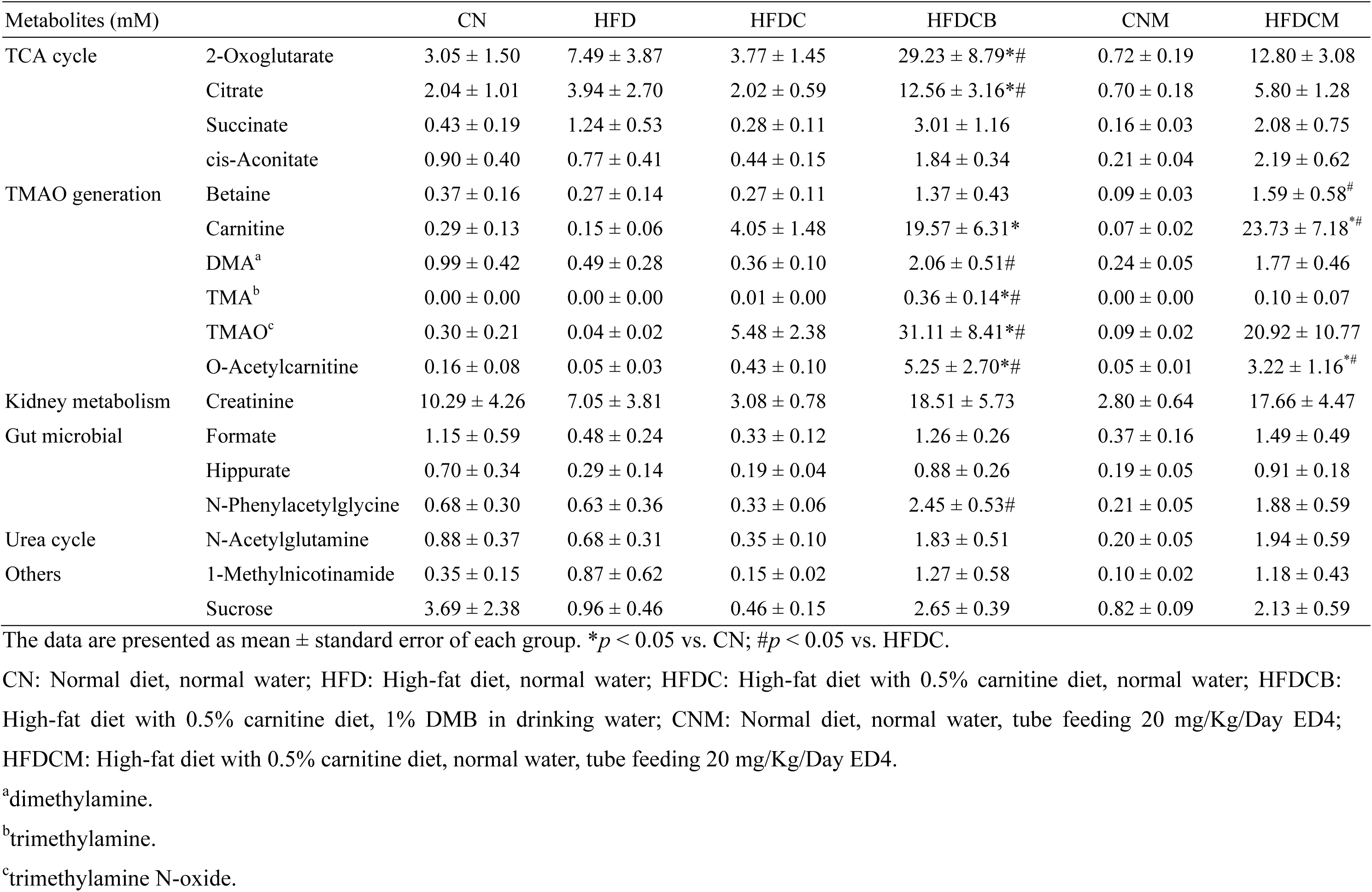
Effects of ED4 on urine metabolite levels in rats with cardiovascular disease induced by high-fat complex carnitine diet.

| Metabolites (mM) |  | CN | HFD | HFDC | HFDCB | CNM | HFDCM |
| --- | --- | --- | --- | --- | --- | --- | --- |
| TCA cycle | 2-Oxoglutarate | 3.05 ± 1.50 | 7.49 ± 3.87 | 3.77 ± 1.45 | 29.23 ± 8.79*# | 0.72 ± 0.19 | 12.80 ± 3.08 |
|  | Citrate | 2.04 ± 1.01 | 3.94 ± 2.70 | 2.02 ± 0.59 | 12.56 ± 3.16*# | 0.70 ± 0.18 | 5.80 ± 1.28 |
|  | Succinate | 0.43 ± 0.19 | 1.24 ± 0.53 | 0.28 ± 0.11 | 3.01 ± 1.16 | 0.16 ± 0.03 | 2.08 ± 0.75 |
|  | cis-Aconitate | 0.90 ± 0.40 | 0.77 ± 0.41 | 0.44 ± 0.15 | 1.84 ± 0.34 | 0.21 ± 0.04 | 2.19 ± 0.62 |
| TMAO generation | Betaine | 0.37 ± 0.16 | 0.27 ± 0.14 | 0.27 ± 0.11 | 1.37 ± 0.43 | 0.09 ± 0.03 | 1.59 ± 0.58<sup>#</sup> |
|  | Carnitine | 0.29 ± 0.13 | 0.15 ± 0.06 | 4.05 ± 1.48 | 19.57 ± 6.31* | 0.07 ± 0.02 | 23.73 ± 7.18*<sup>#</sup> |
|  | DMA<sup>a</sup> | 0.99 ± 0.42 | 0.49 ± 0.28 | 0.36 ± 0.10 | 2.06 ± 0.51<sup>#</sup> | 0.24 ± 0.05 | 1.77 ± 0.46 |
|  | TMA<sup>b</sup> | 0.00 ± 0.00 | 0.00 ± 0.00 | 0.01 ± 0.00 | 0.36 ± 0.14*<sup>#</sup> | 0.00 ± 0.00 | 0.10 ± 0.07 |
|  | TMAO<sup>c</sup> | 0.30 ± 0.21 | 0.04 ± 0.02 | 5.48 ± 2.38 | 31.11 ± 8.41*<sup>#</sup> | 0.09 ± 0.02 | 20.92 ± 10.77 |
|  | O-Acetylcarnitine | 0.16 ± 0.08 | 0.05 ± 0.03 | 0.43 ± 0.10 | 5.25 ± 2.70*<sup>#</sup> | 0.05 ± 0.01 | 3.22 ± 1.16*<sup>#</sup> |
| Kidney metabolism | Creatinine | 10.29 ± 4.26 | 7.05 ± 3.81 | 3.08 ± 0.78 | 18.51 ± 5.73 | 2.80 ± 0.64 | 17.66 ± 4.47 |
| Gut microbial | Formate | 1.15 ± 0.59 | 0.48 ± 0.24 | 0.33 ± 0.12 | 1.26 ± 0.26 | 0.37 ± 0.16 | 1.49 ± 0.49 |
|  | Hippurate | 0.70 ± 0.34 | 0.29 ± 0.14 | 0.19 ± 0.04 | 0.88 ± 0.26 | 0.19 ± 0.05 | 0.91 ± 0.18 |
|  | N-Phenylacetyl glycine | 0.68 ± 0.30 | 0.63 ± 0.36 | 0.33 ± 0.06 | 2.45 ± 0.53<sup>#</sup> | 0.21 ± 0.05 | 1.88 ± 0.59 |
| Urea cycle | N-Acetylglutamine | 0.88 ± 0.37 | 0.68 ± 0.31 | 0.35 ± 0.10 | 1.83 ± 0.51 | 0.20 ± 0.05 | 1.94 ± 0.59 |
| Others | 1-Methylnicotinamide | 0.35 ± 0.15 | 0.87 ± 0.62 | 0.15 ± 0.02 | 1.27 ± 0.58 | 0.10 ± 0.02 | 1.18 ± 0.43 |
|  | Sucrose | 3.69 ± 2.38 | 0.96 ± 0.46 | 0.46 ± 0.15 | 2.65 ± 0.39 | 0.82 ± 0.09 | 2.13 ± 0.59 |

The data are presented as mean ± standard error of each group. \* $p < 0.05$ vs. CN; # $p < 0.05$ vs. HFDC.
CN: Normal diet, normal water; HFD: High-fat diet, normal water; HFDC: High-fat diet with 0.5% carnitine diet, normal water; HFDCB: High-fat diet with 0.5% carnitine diet, 1% DMB in drinking water; CNM: Normal diet, normal water, tube feeding 20 mg/Kg/Day ED4; HFDCM: High-fat diet with 0.5% carnitine diet, normal water, tube feeding 20 mg/Kg/Day ED4.
<sup>a</sup>dimethylamine.
<sup>b</sup>trimethylamine.
<sup>c</sup>trimethylamine N-oxide.

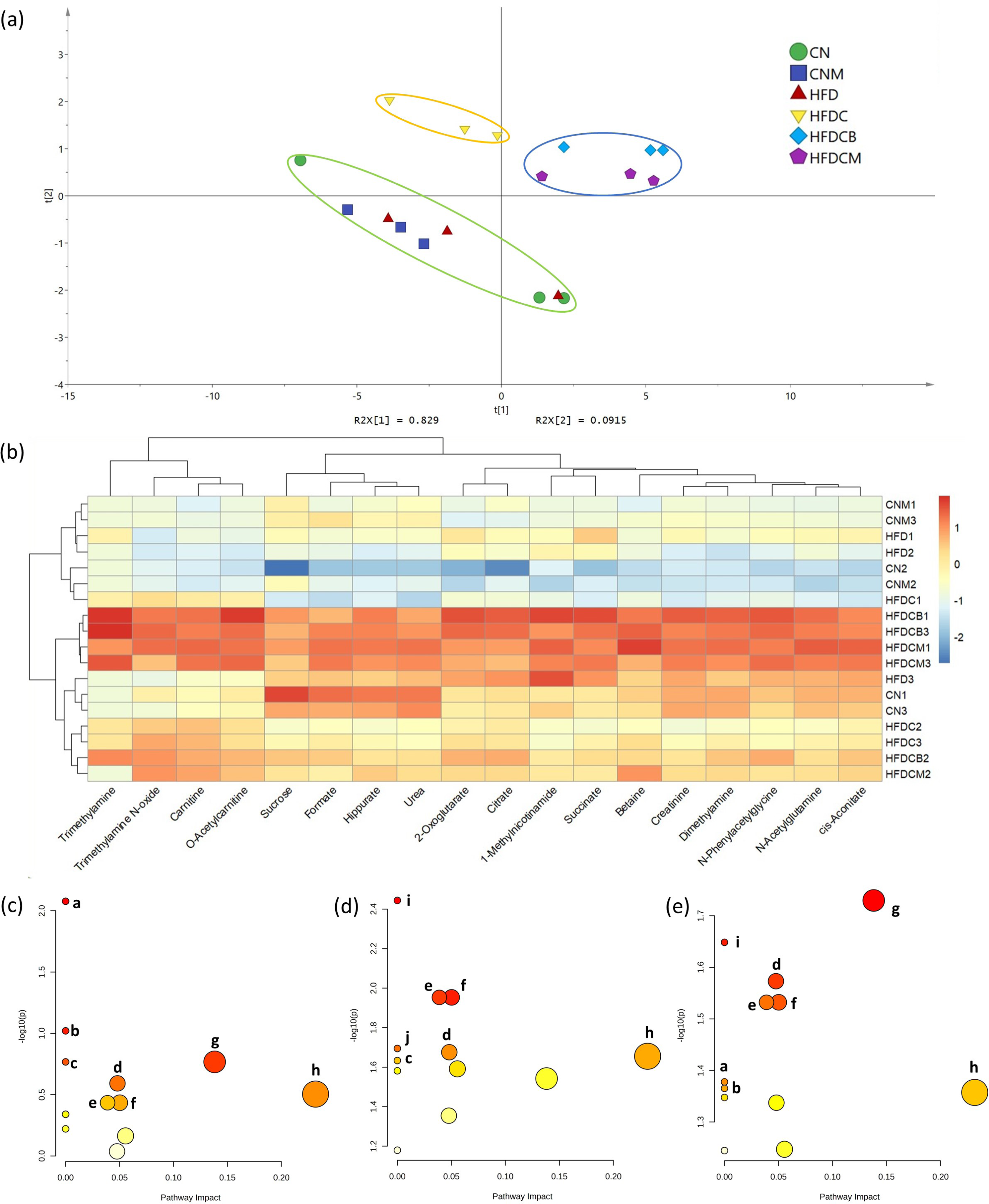

TMA, TMAO, carnitine, and *O*-acetylcarnitine showed a stronger correlation among the urinary metabolites (Fig. 3b). We further compared the levels of carnitine, TMA, and TMAO in the urine across all groups, TMA was not detected in the CN, HFD, and CNM groups (Table 3). However, in the three carnitine-supplemented groups (HFDC, HFDCB, and HFDCM), TMA was detected at concentrations of 0.01, 0.36, and 0.10 mM, respectively. Additionally, TMA levels were higher in the HFDCB and HFDCM groups compared to the HFDC group.

Additionally, carnitine and TMAO levels in the urine of the HFDC, HFDCB, and HFDCM groups were higher than those in the groups not fed carnitine, with the HFDCB and HFDCM groups displaying even greater levels. Specifically, the carnitine levels in the urine of the HFDCB and HFDCM groups were 19.57 and 23.73 mM, representing 4.8- and 5.9-fold increases compared to 4.05 mM in the HFDC group. Similarly, TMAO levels in the HFDCB and HFDCM groups were 31.11 and 20.92 mM, showing 5.7- and 3.8-fold increases compared to 5.48 mM in the HFDC group. However, as shown in Fig. 2, the TMAO levels in the serum of the DMB and ED4 treatment groups were significantly lower than those of the HFDC group. Therefore, we speculate that DMB and ED4 may promote the excretion of TMAO through urine, thereby reducing its concentration in the serum.

The results also showed that the DMB and ED4 groups enhanced the levels of TCA cycle metabolites in urine. Pathway analysis of urinary metabolites, conducted using MetaboAnalyst 6.0 (Fig. 3c-e), revealed that the HFDCB and HFDCM groups influenced several metabolic pathways. Moreover, the results indicated a significant alteration in the TCA cycle for both the DMB and ED4 treated groups, with p-values of 0.0221 and 0.0439, respectively. This suggests that both DMB and ED4 may enhance the TCA cycle and other metabolic pathways, which in turn promotes their excretion.

### FMO3 and OCTN2 Expression

This study analyzed the expression levels of the carnitine transport protein OCTN2 in the jejunum and the TMAO-generating gene FMO3 in the liver (Fig. 4). The OCTN2 expression in the jejunum of the HFDC group was 553.8, which was significantly lower than the 743.8 observed in the CN group. Additionally, FMO3 protein expression in the HFDC group was 1.21, significantly higher than the 0.85 observed in the HFD group. Combined with the elevated TMAO levels in both blood and urine, these findings confirm that carnitine supplementation in the context of a high-fat diet increases FMO3 expression, leading to elevated TMAO production and a corresponding rise in TMAO levels in blood and urine.

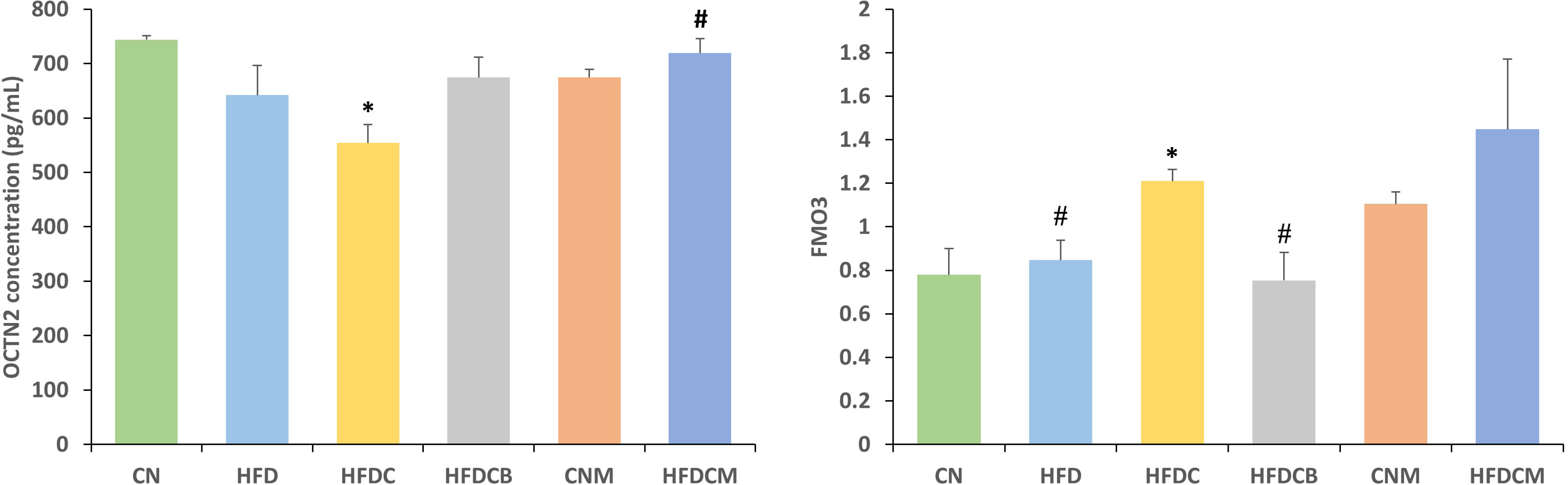

In the HFDCB group, following DMB treatment, the OCTN2 expression level was 674.7, slightly higher than that in the HFDC group, while the FMO3 expression level was 0.75, significantly lower than that in the HFDC group. This suggests that DMB reduces TMAO levels in the blood primarily by inhibiting FMO3 expression.

In contrast, the HFDCM group exhibited a significant increase in OCTN2 expression, reaching 719.3 compared to the HFDC group. However, the FMO3 expression level in the HFDCM group was 1.45, not significantly different from that of the HFDC group. This suggests that the primary mechanism by which ED4 reduces serum TMAO concentrations likely involves enhancing the expression of the carnitine absorption and metabolism gene OCTN2, rather than inhibiting FMO3 expression.

### Gut Microbiota

The *Firmicutes*/*Bacteroidetes* (F/B) ratio results (Fig. 5a) revealed that the F/B ratio in the HFD group was 17.46, significantly higher than the 3.38 observed in the CN group. In the HFDC group, the F/B ratio was 5.72, significantly lower than in the HFD group. The F/B ratio in the ED4-treated HFDCM group was 2.76, which was slightly lower than that of the HFDC group and similar to that of the CN group. Additionally, the F/B ratio in the DMB-treated HFDCB group was 23.91, the highest among all groups, with significant differences compared to both the CN and HFDC groups, indicating that DMB treatment leads to an increase in the F/B ratio.

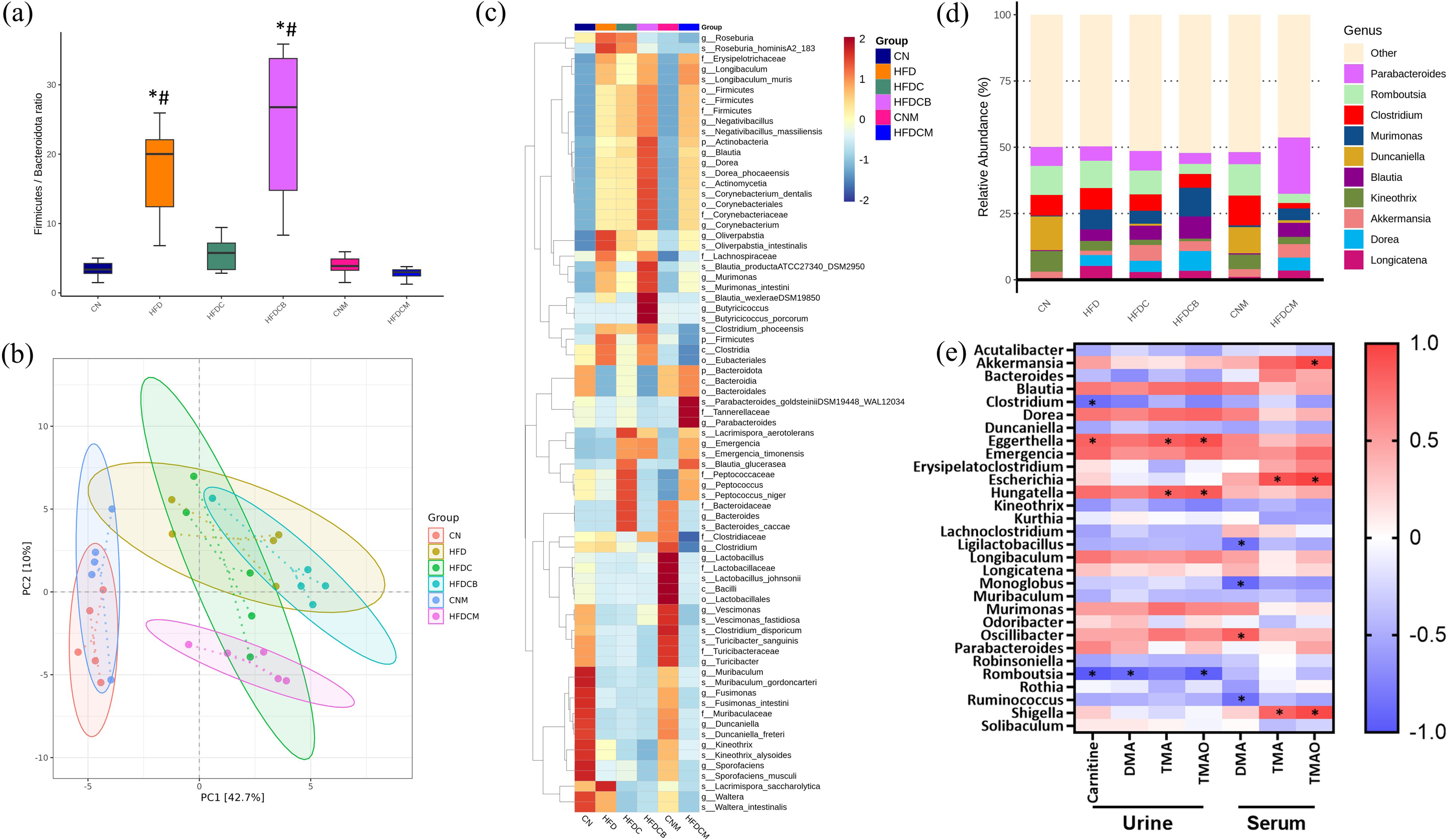

This study analyzed the effects of each treatment group on the intestinal flora of rats. The bacterial composition at the genus level was assessed using PCA (Fig. 5b). The plot shows three distinct clusters: the CN and CNM groups exhibited a high degree of overlap, clearly separated from the other groups, while the HFD, HFDC, and HFDCB groups also showed high overlap. The HFDCM group, however, was separated from the other high-fat diet groups by PC2. Further analysis using linear discriminant analysis (LEfSe) was conducted to identify biomarkers in the feces. A taxonomic cluster heatmap (Fig. 5c), with the LDA score for discriminative features set to 4.0, along with the top ten species abundance chart at the genus level (Fig. 5d), allowed for the comparison of key differential bacterial species between groups. It was found that the relative abundance of *Duncaniella* and *Kineothrix* was higher in the CN and CNM groups compared to the other groups, while the groups fed a high-fat diet (HFD, HFDC, HFDCB, and HFDCM) exhibited significantly higher relative abundances of *Murimonas*, *Blautia*, *Dorea*, and *Longicatena* compared to rats on a normal diet.

Next, we evaluated the effect of carnitine supplementation on the gut microbiota of rats under a high-fat diet. Comparing the relative abundances of the top ten genera between the HFD and HFDC groups, it was found that the HFDC group displayed lower *Murimonas* and *Longicatena,* and higher *Akkermansia* and *Duncaniella*, although these differences were not statistically significant. To further explore the differences between the HFD and HFDC groups beyond the top ten species (Supplementary Fig. S.2), we found that carnitine supplementation significantly increased the relative abundances of *Acutalibacter*, *Bacteroides*, *Eggerthella*, *Emergencia*, *Escherichia*, *Hungatella*, *Muribaculum*, *Robinsoniella*, *Rothia*, and *Shigella*, while significantly decreasing the relative abundances of *Kurthia*, *Odoribacter*, *Monoglobus*, and *Solibaculum*.

To assess whether the treatment groups reversed the bacterial composition observed in the HFDC group, further comparisons were made. In the group treated with DMB, significantly lower relative abundances of *Acutalibacter*, *Bacteroides*, *Escherichia*, *Muribaculum*, *Robinsoniella*, *Rothia*, and *Shigella* were observed, while *Kurthia* and *Odoribacter* showed significantly increased relative abundances (Supplementary Fig. S.3). In the ED4-treated group, a significantly lower abundance of *Bacteroides* and a higher abundance of *Solibaculum* were noted (Supplementary Fig. S.4).

To better understand their correlations, we compared the levels of TMAO and its related metabolites in urine and serum, along with the top 10 genera and microbiota that differed between groups (Fig. 5e). The urinary TMA and TMAO levels were significantly positively correlated with the genera *Eggerthella* and *Hungatella*. Similarly, serum TMA and TMAO levels showed a significant positive correlation with *Escherichia* and *Shigella*. This aligns with the higher abundance of these bacterial genera observed in the high-carnitine group in this study.

## Discussion

RSV has demonstrated beneficial effects on cardiovascular health, while its synthesized derivative, RBE, offers better bioavailability and efficacy than RSV in preventing CVD.^24,32^ Among the monomers of RBE, ED4 not only exhibits the highest partition coefficient but also shows significant potential in combating obesity. Therefore, this study examined the effects of ED4 on TMAO levels, cardiovascular-related indicators, and gut microbiota in a CVD model induced by a high-fat, carnitine-supplemented diet, and explored their interrelationships.

### Recommended HFD with Carnitine Diet Provides More TMAO Than HFD

A high-fat diet is known to disrupt energy homeostasis, leading to overeating and obesity,^33^ while L-carnitine has been shown to reduce body weight, body mass index, and fat mass.^34^ However, excessive supplementation of carnitine can lead to several side effects, including the production of large amounts of TMAO, which may increase the risk of CVD.^35^

This study found that a high-fat diet significantly increased both body weight and body fat in rats without affecting TMAO levels. However, the addition of carnitine to the high-fat diet led to a marked increase in TMAO levels and aortic arch area, along with a slight elevation in serum TG, TC, and AI levels compared to the high-fat diet group alone. Furthermore, administering 1% TMAO in feed has been shown to increase total plasma cholesterol, posing a potential risk for atherosclerosis.^36^ Similarly, other research has found that a high-fat diet combined with carnitine results in a notable increase in atherosclerotic plaque area.^37^ Therefore, we suggest that the TMAO elevation induced by carnitine may cause an increase in TG, TC levels, and aortic arch area. Carnitine has been discussed for its potential role in body weight control and fat reduction.^34^ Consistently, our findings indicate that carnitine supplementation in a high-fat diet can reduce body weight and fat accumulation. However, the HFDC group did not significantly alter the liver fat droplet area compared to the HFD group.

According to the results of this study and existing literature, the combination of a high-fat diet and carnitine supplementation significantly increases serum TMAO levels and shows a tendency to elevate CVD-related indicators. Therefore, a high-fat diet supplemented with carnitine may offer a more suitable model for studying CVD than a high-fat diet alone.

### DMB and ED4 Improve CVD-Related Parameters

DMB, a structural analog of choline, shows potential as a treatment for CVD by inhibiting TMA production by gut microorganisms, thereby reducing TMAO levels.^38^ DMB has been demonstrated to effectively inhibit TMA production in mice fed a high-choline diet.^28^ In this study, we assessed the effect of DMB on TMA production within the context of a high-fat, carnitine-supplemented diet. The results indicated that DMB significantly reduced fat mass, lowered blood pressure, and improved blood lipid levels in rats fed this diet, although its impact on overall body weight was less pronounced. These findings are consistent with previous research, which showed that administering DMB to pregnant rats effectively protected their adult offspring from TCDD-induced hypertension when 1% DMB was added to their drinking water.^39^ Additionally, DMB has been found to inhibit endogenous macrophage foam cell formation and the development of atherosclerotic lesions induced by a choline-rich diet in mice, without altering circulating cholesterol levels.^28^

Our study demonstrated that administering the purified ED4 monomer can mitigate weight gain, body fat, and serum lipid indicators induced by a high-fat, carnitine-supplemented diet. RSV has been shown to lower SBP^40–42^ and reduce the occurrence of hypertension in the offspring of pregnant rats exposed to di-2-ethylhexylphthalate (DEHP)-induced increased blood pressure.^22^ Additionally, it was observed that ED4 reduces body weight and blood lipid levels in rats with HFD-induced obesity.^22,26^ In summary, these findings suggest that both DMB and ED4 treatments have the potential to improve the occurrence of atherosclerosis in rats.

### ED4 and DMB Increase SCFA Levels

SCFAs are the final products of food fermentation by intestinal microorganisms. Acetic acid, propionic acid, and butyric acid are the primary SCFAs in the human gut, accounting for more than 80%. These SCFAs are beneficial to human health, as they can significantly lower plasma triglyceride and cholesterol levels, provide cardiovascular protection, exhibit anti-inflammatory properties, and play a crucial role in maintaining overall health and normal bodily functions.^43,44^

This study analyzed SCFAs in the blood and feces of rats and found that both HFD and HFDC treatments reduced SCFA levels. However, after DMB and ED4 treatments, acetic acid and propionic acid levels were restored to those of the control group. Previous studies have shown that a high-fat diet decreases SCFA levels in the serum of mice, but ED4 treatment can restore SCFA levels.^26^ Other research has also demonstrated that SCFAs play a key role in regulating inflammation, with potential implications for CVD.^45^

Therefore, based on the findings of this study and other research, it can be concluded that ED4 and DMB increase beneficial SCFA levels in rats, which in turn helps reduce hyperlipidemia and provides cardiovascular protection along with anti-inflammatory effects.

### DMB and ED4 May Enhance the Excretion of TMAO in the Urine

The urine metabolomics results observed that the administration of DMB and ED4 following a high-fat, carnitine-supplemented diet increased the levels of various metabolites in the urine. Further comparison of metabolite concentrations in serum and urine showed that the HFDC group had significantly higher serum TMA and TMAO levels than the HFDCB and HFDCM groups. Interestingly, the opposite trend was seen in urine, with the HFDC group having considerably lower TMA and TMAO concentrations compared to the HFDCB and HFDCM groups. Moreover, treatment with DMB and ED4 significantly increased the levels of carnitine and TMAO-related metabolites in urine compared to the HFDC group. These findings suggest that DMB and ED4 may enhance the urinary excretion of TMAO and other metabolites, thereby reducing TMAO levels in the blood.

Some studies have investigated the potential of DMB and ED4 to improve kidney function. DMB has shown promise in managing diabetic kidney disease by inhibiting TMAO levels.^46^ Research indicates that DMB can prevent renal interstitial fibrosis and mitigate impaired renal function caused by a high-fat diet in mice while also reducing circulating TMAO levels.^47^ Limited human studies suggest that RSV administration provides a protective effect against chronic kidney disease, as evidenced by increased kidney filtration rates and volume, along with a mild renal protective effect. Higher doses of resveratrol have also been observed to significantly reduce creatinine levels.^48,49^ It is estimated that nearly 95% of TMA is oxidized to TMAO, which is primarily cleared by the kidneys and excreted in the urine within 24 hours.^50^ However, elevated TMAO levels have been linked to potential adverse effects on kidney health.^51^ Studies have also indicated that kidney function is closely associated with TMAO levels, with serum TMAO concentrations being significantly higher in patients with kidney disease compared to healthy individuals.^52^ These findings underscore the importance of interventions targeting TMAO metabolism as a strategy to improve kidney health and prevent renal dysfunction.

Based on the results of this study and findings from other research, we speculate that the DMB and ED4 treatment groups may enhance the excretion of TMAO derived from dietary carnitine through increased metabolism and elimination via urine, thereby reducing TMAO levels in the blood. However, the mechanisms underlying this action remain unclear and require further investigation.

### Gene

OCTN2 is a carnitine transporter responsible for absorbing carnitine in the intestines and converting it into energy. Once ingested, carnitine is transported to various organs via OCTN2, which is present in both the small and large intestines, providing the body with necessary energy.^53–55^ OCTN2 facilitates the cellular uptake of carnitine and may further contribute to the reduction of TMAO production.^56^ In this study, both DMB and ED4 treatments were shown to enhance OCTN2 expression, with ED4 demonstrating a significantly greater effect compared to HFDC.

Carnitine ingested by the human body is metabolized by the gut microbiota into TMA, which is then absorbed into the bloodstream and ultimately oxidized to TMAO by the FMO3 enzyme in the liver.^57^ Therefore, FMO3 plays an important role in regulating TMAO production. When FMO3 activity is reduced or inhibited, the metabolism rate of TMA decreases, leading to a reduction in TMAO production. A previous study found that plasma TMAO levels in Apoe-/-mice treated with DMB were significantly reduced, along with a marked decrease in total FMO enzyme activity in the liver.^28^ Indeed, in our study, suppression of FMO3 was observed in the DMB-treated group. This indicates that inhibiting the expression of FMO3 is a major mechanism by which DMB reduces TMAO levels.

Our results showed that the ED4 treatment group reduced TMAO levels but exhibited an increase in FMO3 expression. Furthermore, a study investigating the effect of RSV on FMO3 expression and activity in the liver found that RSV treatment significantly increased both FMO3 protein and mRNA levels. This suggests that the reduction in TMAO concentration following RSV treatment is not due to FMO3 regulation in the liver.^58^ Similarly, Bennett et al. reported that overexpression of FMO3 protein in the mouse liver significantly increased plasma TMAO levels.^59^ Therefore, based on the expression of TMAO-related genes, we speculate that the regulation of OCTN2 may be one of the factors contributing to the reduction of TMAO levels by ED4, whereas FMO3 is not.

### The Effect of Gut Microbiota

There is an inseparable relationship between gut microbiota and TMA production. Therefore, improving gut microbiota presents a rapid and effective approach to reducing TMAO levels in the blood. However, the relationship between gut bacteria and metabolites is highly complex and subject to individual variation. Differences in experimental methods and TMA induction modes contribute to the diversity of TMA-producing strains reported in various studies.^60–63^

This study also evaluated the impact of different treatment groups on intestinal bacteria. Significant differences in the relative abundance of several bacterial genera were found between the HFD and HFDC groups. Additionally, the addition of carnitine to the diet significantly increased the abundance of microbes such as *Acutalibacter*, *Bacteroides*, *Emergencia*, *Escherichia*, *Hungatella*, *Muribaculum*, *Robinsoniella*, and *Shigella*, all of which have been discussed in other literature in relation to TMA production. The increased abundance of *Acutalibacter*, *Bacteroides*, *Muribaculum*, and *Robinsoniella* following carnitine supplementation has also been confirmed to correlate with higher TMAO levels in the body.^64–67^ Studies have pointed out that *Emergencia* and *Escherichia* spp. possess TMA-producing genes capable of metabolizing carnitine to produce TMA,^62,68^ and *Shigella* species have also been identified as important TMA producers.^13^ The expression level of the TMA-producing gene was also found to be positively correlated with *Hungatella*.^69^ Moreover, *Escherichia*, *Shigella*, and *Hungatella* also showed a positive correlation with TMA and TMAO levels in either urine or serum in our study.

Contrary to some of the findings in this study, other sources suggest that *Kurthia* is a TMA-producing bacterium,^70^ and *Odoribacter* is highly prevalent in individuals with high TMAO concentrations in healthy gut microbiota.^67^ However, in this study, the relative abundance of these two bacteria was significantly lower in the carnitine diet group compared to the high-fat diet group.

The DMB-treated group showed a reduction in the relative abundance of TMA-producing bacteria, including *Acutalibacter*, *Bacteroides*, *Muribaculum*, *Robinsoniella*, and *Shigella*, demonstrating that DMB can alter the gut microbiota influenced by a high-carnitine diet.

In the ED4-treated group, the effect of inhibiting TMA-producing bacteria was not as significate as in the DMB group. However, we still observed the ED4 treatment group significantly reduced the relative abundance of Bacteroides. These results indicate that ED4 can improve certain gut microbiota changes caused by carnitine, although it is not as effective as DMB.

Additionally, a higher F/B ratio is often used as an indicator of obesity and is also considered to be associated with CVD and other diseases.^71^ In this study, the HFD group had a higher F/B ratio, accompanied by greater weight gain, visceral fat, and peripheral fat. Moreover, the HFDC and HFDCM groups exhibited a significantly lower F/B ratio than the HFD group, along with reduced weight gain, fat mass, and fat accumulation. Interestingly, while the DMB group displayed the highest F/B ratio among all groups, it also showed significant inhibition of weight gain, fat mass, and fat accumulation.

Magne et al. reviewed the correlation between the F/B ratio and obesity, concluding that many factors influence this ratio, making it challenging to use as a definitive hallmark of obesity.^71^ Based on our results, we partially agree with their conclusion. However, we suggest that the F/B ratio can still serve as a reference indicator in studies on obesity and related diseases.

### Conclusion

This study established an innovative animal model using a high-fat diet supplemented with carnitine to induce TMAO elevation, effectively mimicking dietary-induced cardiovascular risk factors (Fig. 6). This model offers a reliable experimental platform for exploring potential therapeutic strategies for cardiovascular disease (CVD). The findings highlight the therapeutic potential of resveratrol butyrate ester (ED4) in mitigating CVD and related metabolic disorders. ED4 supplementation effectively lowers blood lipid levels, reduces fat droplet accumulation, and decreases serum TMAO levels, collectively suggesting a reduced risk of atherosclerosis. ED4’s multimechanistic effects likely involve the upregulation of the carnitine absorption gene OCTN2, promoting carnitine utilization and enhancing the urinary excretion of TMAO and its related metabolites. Moreover, ED4 improves gut microbiota composition, notably reducing the abundance of TMA-producing Bacteroides and restoring short-chain fatty acid (SCFA) concentrations to near-normal levels. These findings suggest that ED4 reduces cardiovascular risk factors by modulating gut microbiota and metabolic pathways, ultimately lowering bloodstream TMAO levels. This study contributes to the field by introducing the first animal model combining a high-fat diet and carnitine supplementation to simulate diet-induced cardiovascular risk mechanisms, while also demonstrating ED4’s potential in anti-obesity, metabolic regulation, and cardiovascular protection. However, while the study provides evidence of ED4’s ability to enhance TMAO excretion via urine, the underlying molecular mechanisms remain unclear due to limited supporting gene expression data. Future research should focus on elucidating these mechanisms, particularly the regulatory pathways involved in TMAO metabolism, to comprehensively understand ED4’s role in cardiovascular health and its potential translational applications.

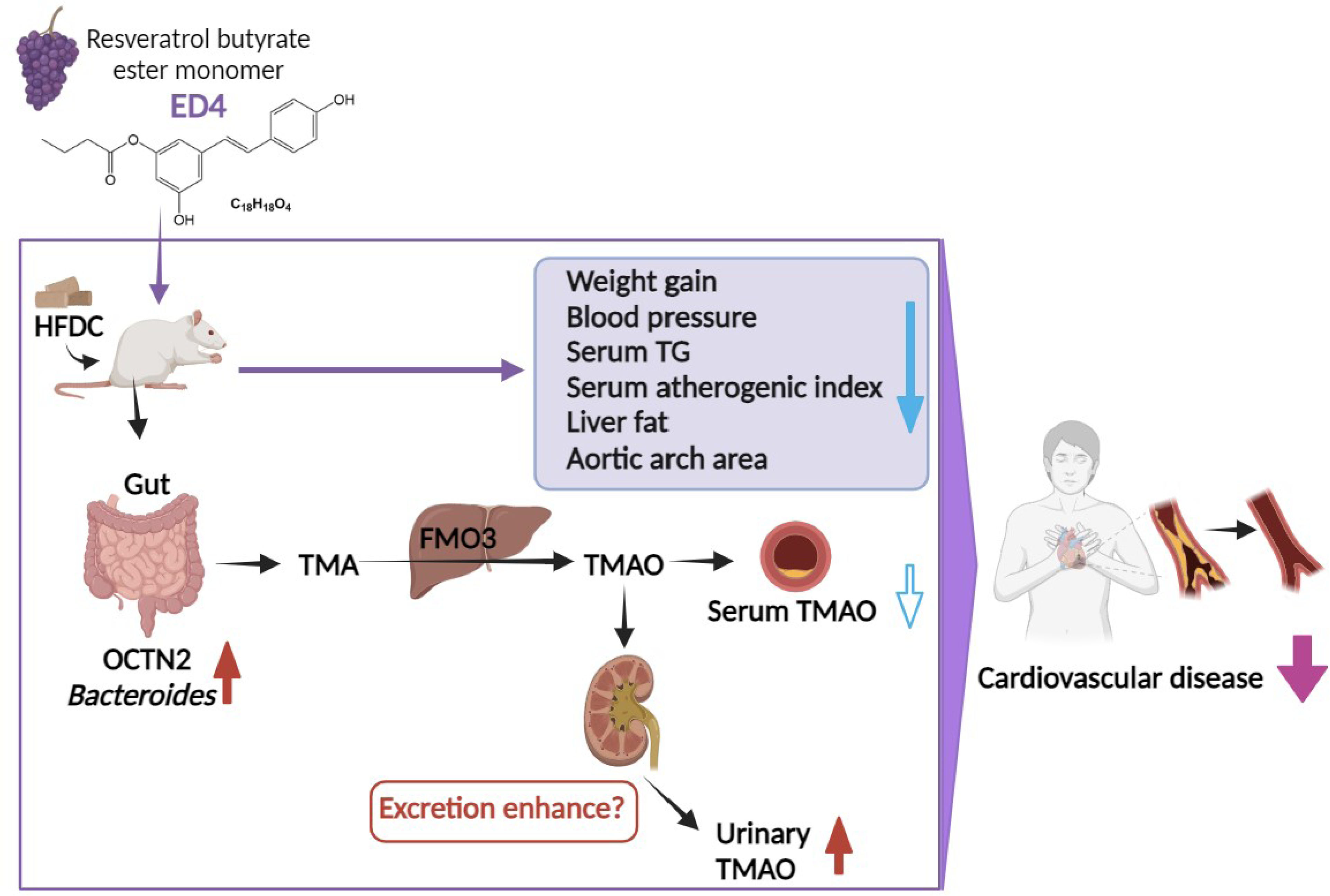

## CRediT authorship contribution statement

**Chih-Yao Hou:** Conceptualization, data curation, investigation, project administration, supervision, writing—review and editing. **Cai-Sian Liu:** Conceptualization, data curation, formal analysis, investigation, software, validation, visualization. **Ming-Kuei Shih:** Conceptualization, investigation, project administration, writing—review and editing. **Asif Ali Bhat:** Investigation, writing—original draft. **You-Lin Tain:** Conceptualization, project administration, supervision. **Chang-Wei Hsieh:** Conceptualization, supervision. **Yu-Wei Chen:** Conceptualization, data curation, formal analysis, investigation, software, validation, writing—review and editing. **Shin-Yu Chen:** Project administration, software, validation, visualization, writing—original draft, writing—review and editing.

## Data availability

Data will be made available on request.

## Acknowledgments

The authors would like to acknowledge all the individuals who volunteered for this study.

## Appendix A. Supplementary data

The following are the Supplementary data to this article: Supplementary data 1.

